# GLIS3 is a key regulator of astrocyte differentiation in human neural stem cells

**DOI:** 10.64898/2026.04.02.716227

**Authors:** Tapas Pradhan, Hong Soon Kang, Kilsoo Jeon, Sara A. Grimm, Kye-yoon Park, Anton M. Jetten

## Abstract

Astrocytes play a key role in neuronal homeostasis and in various neural disorders. The generation of astrocytes from neural progenitor cells (NPCs) and its functions are under a complex control of several signaling networks and transcription factors. In this study, we demonstrate that the transcription factor, GLIS similar 3 (GLIS3), which has been implicated in several neurodegenerative diseases, is highly expressed in astrocytes, and is required for the efficient differentiation of human NPCs into astrocytes. Loss of GLIS3 function greatly impairs astrocytes differentiation, resulting in reduced expression of astrocyte markers, whereas expression of exogenous GLIS3 restores the induction of astrocyte specific genes indicating a critical role for GLIS3 in astrocyte differentiation. Integrated transcriptomic and cistromic analyses revealed that GLIS3 directly regulates the transcription of several astrocyte-associated genes, including *GFAP*, *SLC1A2*, *NFIA*, and *ATF3*, in coordination with lineage-determining factors, such as STAT3, NFIA, and SOX9. We hypothesize that GLIS3 dysfunction disrupts this transcriptional network thereby contributing to astrocyte-associated neurological disorders. Identification of GLIS3 as a key regulator of astrocyte differentiation and gene expression will advance our understanding of its role in neurodegenerative diseases and may provide a new therapeutic target.

## Introduction

GLI-similar 3 (GLIS3) is a Krüppel-like zinc-finger transcription factor that regulates gene expression by binding G-rich GLIS binding sites (GLISBS) within the regulatory regions of target genes and either activates or represses their transcription depending on the cellular and promoter context (Kim et al. 2003; Jeon et al. 2019; Scoville et al. 2019; Kang et al. 2023). Loss of GLIS3 function causes dysregulation of gene expression and impairment of various biological processes in several tissues leading to multiple pathologies, including diabetes, polycystic kidney disease, and hypothyroidism (Kim et al. 2003; Kang et al. 2009; Kang et al. 2017; Scoville et al. 2020). There is growing evidence for a role of GLIS3 in the human nervous system and neurological disorders (Senee et al. 2006; Jeon et al. 2019; Dimitri, 2017 #3}. Genome wide association studies (GWAS) have identified an association between genomic variants of *GLIS3* and increased risk of Alzheimer’s (AD) and Parkinson’s (PD) disease (Cruchaga et al. 2013; Chung et al. 2015a; Calderari et al. 2018; Insel et al. 2023). In addition, *GLIS3* has been reported to be dysregulated in AD (Hill and Gammie 2022) and a correlation was found between *GLIS3* variants and elevated levels of total tau and phosphorylated tau in cerebrospinal fluid (Cruchaga et al. 2013). GLIS3 has further been shown to promote the differentiation of human embryonic stem cells (hESCs) into posterior neural progenitor cells (NPCs) in lieu of the default anterior pathway by enhancing the expression of the posteriorizing factor WNT3A (Jeon et al. 2019).

In this study, we demonstrate that GLIS3 is highly expressed in astrocytes suggesting a regulatory function in these cells. Astrocytes are the most abundant glial cells and are important in maintaining the health and function of the central nervous system (CNS), including regulation of the extracellular ion balance, synapse formation, maintaining blood-brain barrier function, roles in injury repair mechanisms and neuroinflammation (Christopherson et al. 2005; Allen et al. 2012; Dienel 2019; Durkee and Araque 2019; Zhou et al. 2019; Jackson et al. 2022; Valles et al. 2023). They play an active role in synaptic transmission by regulating the uptake and release of neurotransmitters and influencing neuronal communication (Chung et al. 2015b; Cuellar-Santoyo et al. 2022). Dysregulation of astrocyte function has been implicated in multiple neurodegenerative diseases, including AD, PD, and Huntington’s disease (Tyzack et al. 2016; Siracusa et al. 2019; Brandebura et al. 2023).

During CNS development, astrocytes arise from neural progenitor cells (NPCs) through a temporally regulated process known as astrogliogenesis (Ferreira and Pinto 2025). The gliogenic switch, the transition from neuronal fate of NPCs to astrocytes, is a tightly regulated process orchestrated by signaling cascades, including the Notch pathway, activation of JAK/STAT by bone morphogenic proteins (BMPs), and regulation by several transcription factors, such as SOX9, NFIA, and STAT3 (Sloan and Barres 2014; Garg et al. 2025; Jin et al. 2025). Together these pathways suppress neurogenesis and promote astrogliogenesis and the expression of astrocyte-specific gene expression, such as glial fibrillary acidic protein (GFAP) and the calcium binding protein S100B (Ma et al. 2024). While important insights have been obtained into the regulation of murine gliogenesis, the regulatory processes governing human astrocyte lineage specification is not fully understood. Differentiation of human embryonic stem cells (hESCs) into different neural cell types, including astrocytes, provides a powerful experimental model to study the regulatory networks underlying human astrogliogenesis (Chandrasekaran et al. 2016; Prajumwongs et al. 2016; Perriot et al. 2021).

To obtain insights into the role of GLIS3 in astrogliogenesis, we analyzed the effect of loss-of-GLIS3 function on the stepwise differentiation of hESCs into astrocytes by transcriptomics and cistromics using wild type (WT) and GLIS3-deficient hESCs as experimental models. Our study identifies GLIS3 as a critical transcriptional regulator of astrocyte differentiation and gene expression and expands our knowledge of GLIS3 function in the nervous system and will enhance our understanding of its role in neurodegenerative diseases, thereby inspiring the development of future therapeutic strategies.

## Results

### GLIS3 is highly expressed in astrocytes

Analysis of the brain RNA-seq data base (www.brainrnaseq.org) indicated that *Glis3* was highly expressed in both human and murine astrocytes compared to several other neural cell types (Fig. 1A). This was confirmed by qPCR analysis showing that the level of *Glis3* mRNA expression in isolated mouse astrocytes was comparable to that in kidney, a tissue, in which *Glis3* is known to be highly expressed (Fig. 1B; Supplemental Fig. S2A). Analysis of GLIS3 subcellular localization in cultured astrocytes isolated from PND3-4 *Glis3*-GFP mice, expressing a GLIS3-GFP fusion protein, showed that GLIS3-GFP was largely restricted to the nucleus of GFAP^+^ astrocytes, whereas F4/80^+^ microglia showed very weak staining (Fig. 1C; Supplemental Fig. S2B).

**Figure 1.**
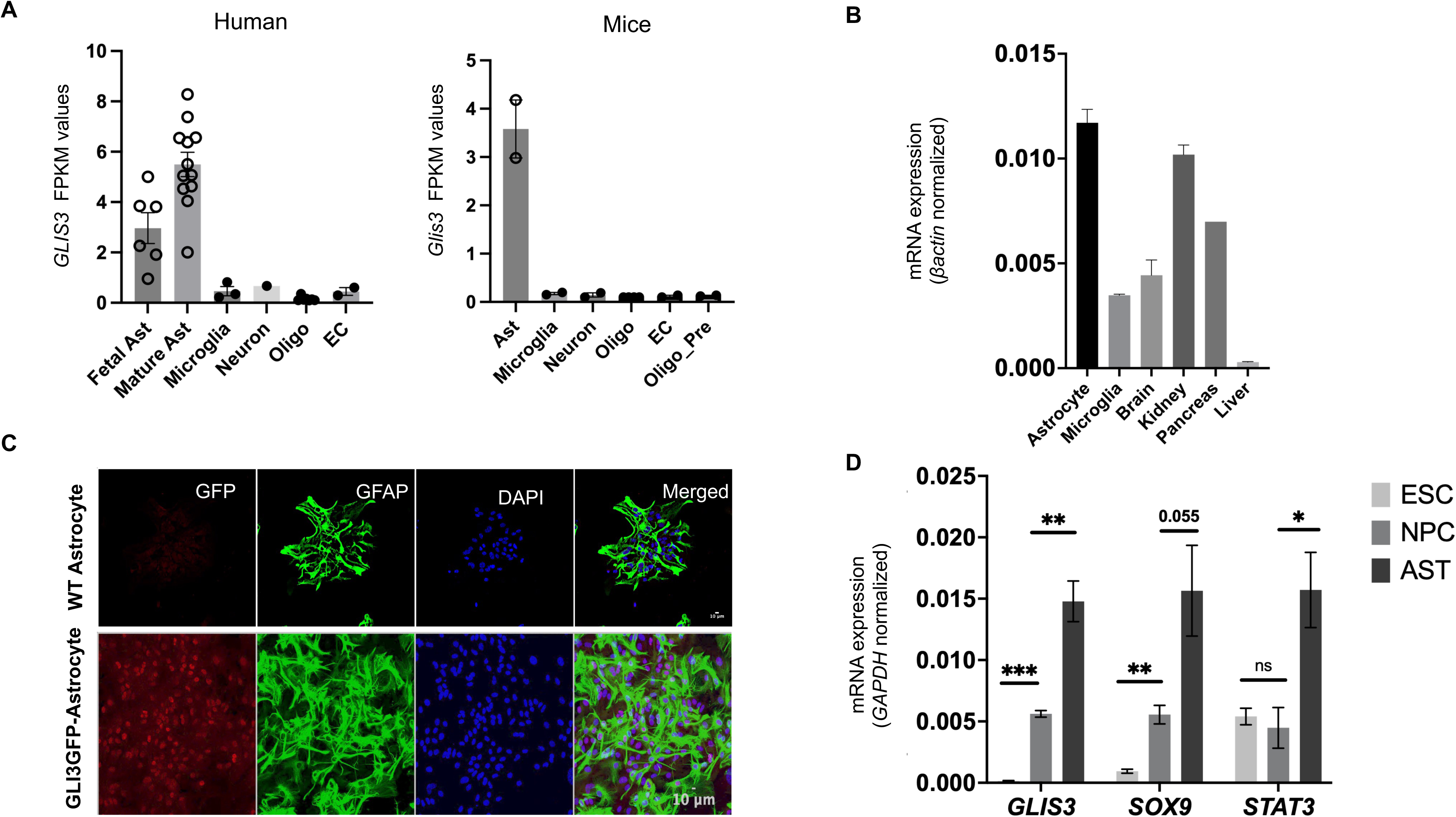
GLIS3 is highly expressed in astrocytes. (A) GLIS3 expression in different cell types of the human and mouse CNS obtained from the RNA seq database, www.brainrnaseq.org. Abbreviations: Ast, Astrocytes; Oligo, Oligodendrocytes; Oligo_Pre, Oligodendrocyte precursor; EC, Endothelial cells. (B) qPCR analysis of Glis3 expression in several mouse-derived cells and tissues. (C) Co-immunostaining of GLIS3-GFP and GFAP in primary astrocytes. Scale bar 10 µm. (D) Expression of GLIS3, SOX9 and STAT3 mRNA at different stages during the differentiation hESCs into astrocytes (n=3). Abbreviations: ESC, embryonic stem cells; NPC, neural progenitor cells; AST, astrocytes. The data are presented as SEM values, with ^∗^ p<0.05; ^∗∗^ p<0.01; ^∗∗∗^ p<0.001; ns, not significant.

Next, we examined whether *GLIS3* expression was temporally regulated during the differentiation of hESCs into NPCs and their subsequent differentiation into astrocytes. As shown in Fig. 1D, GLIS3 mRNA expression was very low in hESCs and induced upon their differentiation into NPCs and increased further upon their differentiation into astrocytes. The pattern of *GLIS3* expression was comparable to that of *SOX9*, encoding a transcription factor with a critical regulatory role in NPCs and astrocytes (Fig. 1D). Its high expression in astrocytes suggested a role for GLIS3 in the regulation of astrocyte differentiation or function.

### GLIS3 is required for the differentiation of hNPCs into astrocytes

To obtain insights into the role of GLIS3 in astrocyte differentiation and/or function, we examined the effect of loss of GLIS3 function on the differentiation of hESCs into NPCs and subsequently into astrocytes using WT and *GLIS3*^-/-^ hESCs. To assess potential clonal heterogeneity, we used two independent *GLIS3*^-/-^ clones that yielded similar results. GLIS3-deficiency had no significant effect on ESC morphology, while the level of expression of the pluripotent markers, *OCT4, NANOG,* and *SOX2,* as well as the immunostaining of the respective proteins were similar between WT and *GLIS3*^-/-^ cells (Fig. 2 A, B; Supplemental Fig. S3A-C). The pluripotency of *GLIS3*^-/-^ ESCs was supported by their ability to differentiate into all three germ layers as shown by the expression OTX2, SOX17 and TBXT that was comparable to that of WT hESCs (Supplemental Fig. S3D). These results indicated that GLIS3 does not play a major role in the maintenance or pluripotency of hESCs, consistent with *GLIS3*’s minimal expression in hESCs.

**Figure 2.**
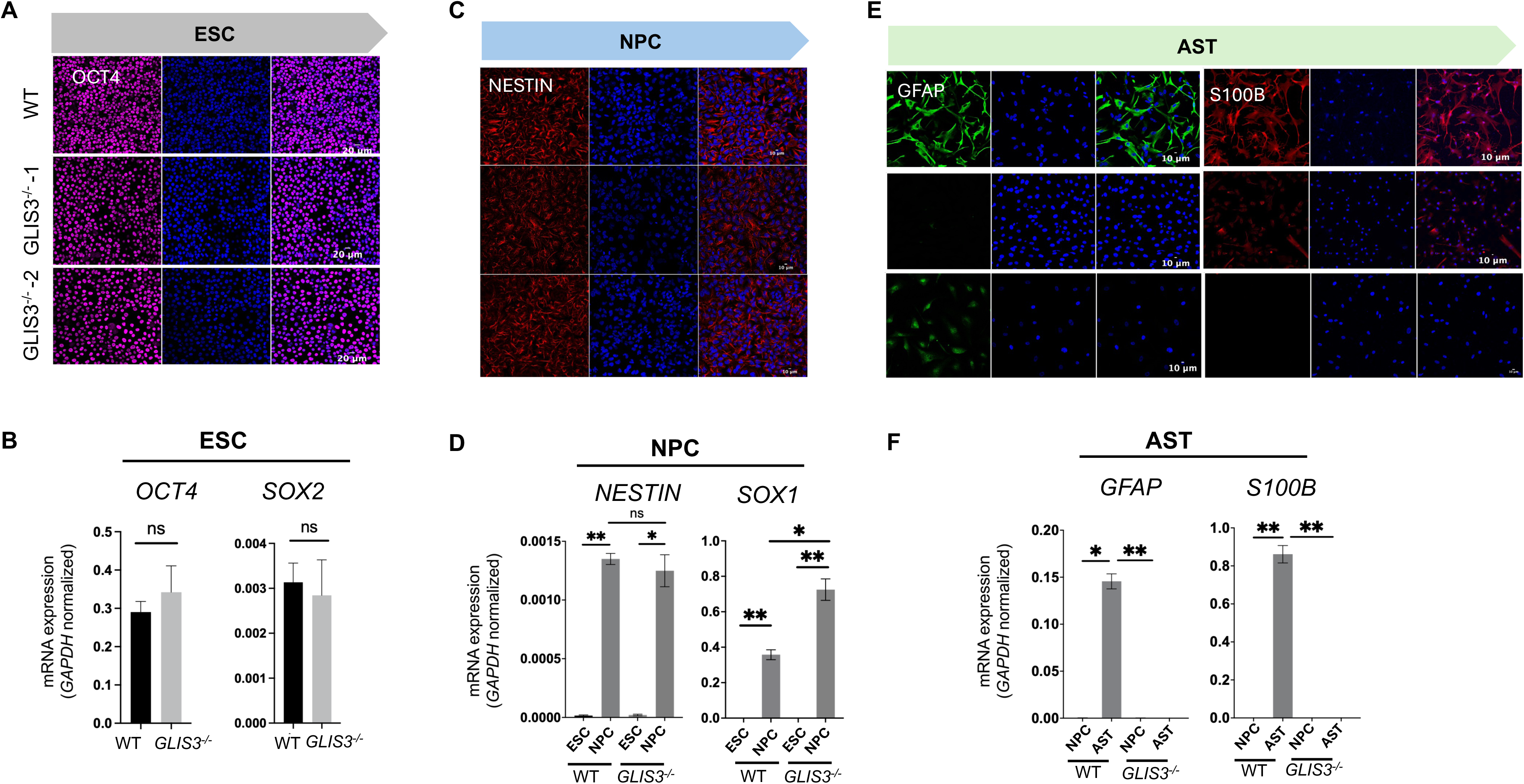
GLIS3 is required for the differentiation of hNPCs into astrocytes. (A) Representative immunofluorescence images showing expression and nuclear localization of OCT4 protein in WT and *GLIS3*^-/-^ hESCs. Scale bar 20 µm (B) Bar graph showing qPCR quantification of *OCT4* and *SOX2* expression in WT and *GLIS3*^-/-^ hESCs (n=3). (C) Representative immunofluorescence images showing expression and localization of NESTIN in WT and *GLIS3*^-/-^ NPCs. Scale bar 10 µm (D) qPCR analysis of *NESTIN* and *SOX1* expression in WT and *GLIS3*^-/-^ NPCs (n=3). (E) Representative immunofluorescence images showing expression and localization of GFAP and S100B protein in WT and GLIS3^-/-^ NPCs induced to differentiate into astrocytes. Scale bar 10 µm. (F) qPCR analysis of *GFAP* and *S100B* expression in WT and *GLIS3*^-/-^ NPCs induced to differentiate into astrocytes (n=3). The data are presented as SEM values, with ^∗^ p<0.05; ^∗∗^ p<0.01; ns, not significant.

Next, we examined the differentiation of hESCs into NPCs. After one week of differentiation of hESCs into NPCs, both WT and *GLIS3*^-/-^ cells underwent the typical changes in cellular morphology from small, compact cells to larger, epithelial-like cells (Supplemental Fig. S3E). qPCR analysis showed that the differentiation of both WT and *GLIS3*^-/-^ hESCs into NPCs was accompanied by decreased expression of *OCT4* and increased expression of the NPC marker genes*, NESTIN and SOX1.* No significant differences were observed in the level of *NESTIN* mRNA expression between WT and *GLIS3*^-/-^ NPCs, while the fold increase in *SOX1* expression was slightly higher in *GLIS3*^-/-^ NPCs compared to WT NPCs. NPC differentiation was further supported by immunostaining for NESTIN and SOX1; no significant difference was observed between WT and *GLIS3*^-/-^ NPCs. (Fig. 2C, D; Supplemental Fig. S3F, G), Subsequently, we examined whether loss of GLIS3 function affected the ability of NPCs to differentiate into astrocytes. Differentiation of WT NPCs into astrocytes was accompanied by induction of the astrocyte markers, *GFAP* and *S100B* (Supplemental Fig. S4); however, these genes were not induced in *GLIS3*^-/-^ cells (Fig. 2E). The observations were supported by immunostaining, which detected GFAP and S100B in WT astrocytes, but not in *GLIS3*^-/-^ cells (Fig. 2F). These findings suggested that GLIS3 function is required for proper differentiation of hNPCs into astrocytes.

### Regulation of gene expression by GLIS3 during NPC-astrocyte differentiation

To obtain further insights into the effects of GLIS3 deficiency on the global changes in gene expression during the differentiation of NPCs into astrocytes, we performed RNA-seq analysis (Supplemental Fig. S5A). Comparison of the gene expression profiles between WT astrocytes and *GLIS3*^-/-^ cells showed that of the differentially expressed genes (DEGs; FDR <0.01), 1805 genes were upregulated, while 2061 genes were suppressed in *GLIS3*^-/-^ cells (Fig. 3A). All DEGs and associated pathways are listed in Supplemental Tables S5-8. DEG analysis with astrocyte-related GSEA gene sets identified astrocyte development, differentiation, and astrocyte markers as negatively enriched processes in *GLIS3*^-/-^ cells (Fig. 3B). This was supported by Gene Ontology (GO) analysis showing that genes involved in neural system development and astrocyte function, such as axon guidance, synaptic transmission formation, and ionotropic glutamate signaling, were among the top processes suppressed in *GLIS3*^-/-^cells (Fig. 3C). This included *GFAP, S100B, FABP7, SLC1A2 (GLT1* or *EAAT2*), *ALDH1L1*, *SLIT1*/2, *NLGN1*, and *GRIA4* (Fig. 3D), the GABA transporters SLC6A11. In addition, the expression of several genes encoding TFs with critical roles in the regulation of astrocyte differentiation and function, including *NFIA, SOX9, SOX2, SOX8, ATF3, RORB, and LHX2*, was suppressed in *GLIS3*^-/-^ cells (Fig. 3D). The repression of several of these genes was supported by qPCR analysis (Fig. 3E). Together, these data indicated that loss of GLIS3 function inhibits the differentiation of NPCs into astrocytes and suggested that this may be in part related to the suppression of several TFs that are critical for astrocyte differentiation.

**Figure 3.**
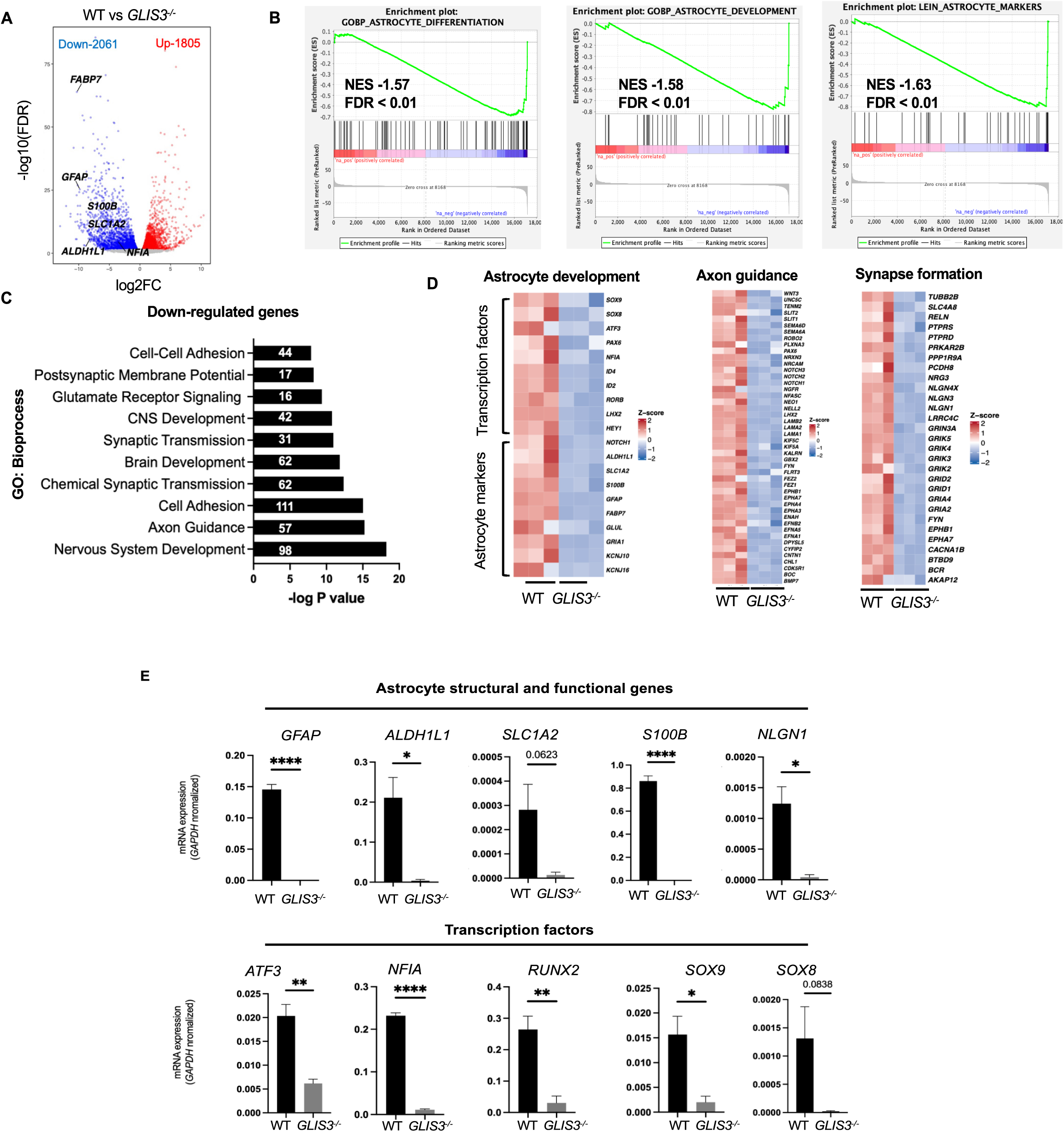
Regulation of gene expression by GLIS3 during NPC-astrocyte differentiation. (A) Volcano plot showing genes differentially expressed (DEGs) in *GLIS3*^-/-^ cells. The X-axis showing the log2-fold change in gene expression and the Y-axis the log10 P-value. Blue, downregulated genes; Red, up-regulated genes; Grey, non-significant change at FDR 0.01. (B) Gene set enrichment analysis (GSEA) of DEGs from *GLIS3*^-/-^ cells with publicly available human astrocyte gene sets. (C) Gene Ontology analysis of genes suppressed in *GLIS3*^-/-^ cells showing enrichment in various bioprocesses. (D) Heatmaps of genes involved in, respectively, astrocyte development, axon guidance and synapse formation. (E) qPCR analysis of several genes associated with astrocyte development and function in WT and *GLIS*3^-/-^ NPCs induced to differentiate into astrocytes (n=3). The data are presented as SEM values, with ^∗^ p<0.05; ^∗∗^ p<0.01; ^∗∗∗∗^ p<0.0001.

Pathway analysis of genes expressed at a higher level in *GLIS3*^-/-^ cells compared to WT cells identified cell division, cell cycle, and mitosis as the top pathways (Supplemental Fig. S5B). The increase in the expression of various cell cycle-related genes, including several cyclins, is shown in the heatmap in Supplemental Fig. S5C. These observations are consistent with the concept of an inverse relationship between cell proliferation and differentiation observed in many cell systems, including the differentiation of NPCs into astrocytes (Ruijtenberg and van den Heuvel 2016).

### GLIS3 directly regulates the transcription of astrocyte-associated genes

GLIS3 regulates transcription by binding to a GC-rich consensus sequence (GLISBS) in the regulatory regions of target genes (Scoville et al. 2020; Kang et al. 2023). To determine which genes are directly regulated by GLIS3 in astrocytes, we performed ChIP-seq analysis with astrocytes derived from hESCs expressing GLIS3-HA. The peak calling algorithm identified a total of 66,687 GLIS3 binding peaks, of which 44.46% and 31.86% were localized within the gene body and intergenic regions, respectively, and 11.46% and 12.22% were associated with, respectively, the upstream and proximal promoter regions (Fig. 4A-C). HOMER analysis identified a GLIS3 consensus-binding site among the top sequences, as well as consensus binding sites for SOX, NFI, STAT, and AP-1 transcription factors (Fig. 4D). Members of these families, including SOX9, SOX2, STAT3, cJUN, and NFIA/B, play a critical role in the regulation of astrocyte proliferation and differentiation, and the expression of astrocyte marker genes, including *GFAP, SLC1A2,* and *S100B* (Bachetti et al. 2010; Saito et al. 2018; Pavlou et al. 2019; Han et al. 2020). These observations support the hypothesis that GLIS3 regulates astrocyte gene transcription and functions in coordination with other TFs, such as SOX9, STAT3, NFIA, and AP-1.

**Figure 4.**
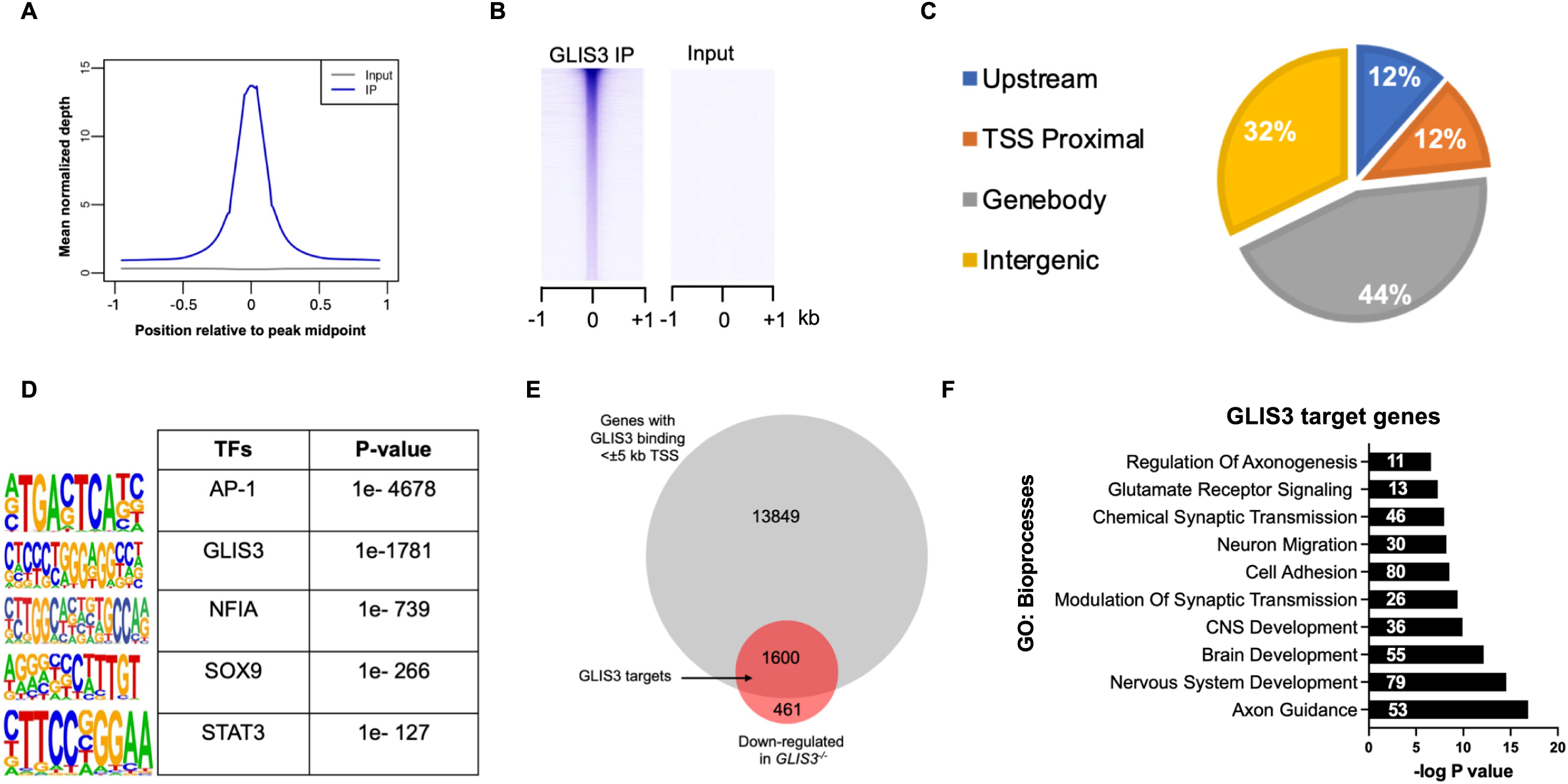
GLIS3 directly regulates the transcription of astrocyte-associated genes. (A-B) ChIP-Seq read density and heatmap of the 2 kb region centered on each of the GLIS3 binding peaks identified. (C) Binding analysis of GLIS3 ChIP-Seq data showing preference of GLIS3 binding across the genome. (D) HOMER analysis of ChIP-Seq data showing binding motifs of GLIS3 and other neighboring motifs protein near GLIS3 binding regions. (E) Venn diagram showing overlap between GLIS3 bound genes (<±5 kb of TSS) and down-regulated genes obtained from ChIP-seq and RNA-seq analysis, respectively. (F) GO enrichment analysis of direct targets genes of GLIS3 in astrocytes.

In determining which DEGs were directly regulated by GLIS3, we limited our analysis to the 15450 genes with a GLIS3 binding peak within <± 5 kb of transcriptional start site (TSS). As shown in the Venn diagram in Fig. 4E, 77.6% (1600 genes) of the suppressed genes and 68.3% (1233 genes) of the genes upregulated in *GLIS3*^-/-^ cells were GLIS3 targets (Fig. 4E; Supplemental Fig. S6A). These data our consistent with our concept that GLIS3 can function both as a transcriptional activator as well as a repressor (Kim et al. 2003; Jeon et al. 2019). GO pathway analysis of the suppressed target gene set identified CNS development, axon guidance, and synapse formation among the top processes, while cell division and cell cycle were associated with the target genes that were higher expressed in *GLIS3*^-/-^ cells (Fig. 4F; Supplemental Fig. S6B). These pathways are very similar to those identified by GO analysis of all DEGs (Fig. 3C; Supplemental Fig. S5B). Supplemental Tables S9-13 show lists of GLIS3 target genes and associated pathways. The suppression of GLIS3 target genes with an established role in astrocyte development and function (Table 1) agrees with a regulatory role for GLIS3 in astrocytes.

**Table. 1.** List of astrocyte specific genes bound and regulated by GLIS3.

| Genes | log2 FC | FDR | GLIS3 ChIP-seq signal |
| --- | --- | --- | --- |
| *FABP7* | -10.3 | 1.31E-64 | Yes |
| *GFAP* | -9.8 | 4.91E-26 | Yes |
| *ID4* | -8.8 | 5.53E-25 | Yes |
| *ALDH1L1* | -8.1 | 1.09E-06 | Yes |
| *S100B* | -7.1 | 1.21E-22 | Yes |
| *SOX8* | -5.1 | 0.000105169 | Yes |
| *HEY1* | -4.9 | 1.86E-18 | Yes |
| *SLC1A2* | -4.6 | 2.29E-09 | Yes |
| *ID2* | -3.7 | 4.95E-13 | Yes |
| *NFIA* | -3.6 | 1.68E-05 | Yes |
| *SOX9* | -1.8 | 1.36E-05 | Yes |
| *ATF3* | -1.8 | 0.002687575 | Yes |
| *GRIA2* | -8.0 | 2.12E-30 | Yes |
| *NLGN3* | -7.5 | 7.64E-26 | Yes |
| *SEMA6A* | -7.2 | 1.79E-24 | Yes |
| *SLIT1* | -6.7 | 5.70E-07 | Yes |
| *NLGN1* | -6.3 | 1.74E-30 | Yes |
| *ROBO2* | -5.3 | 2.46E-14 | Yes |
| *RELN* | -5.0 | 2.31E-05 | Yes |
| *EPHB1* | -4.9 | 3.93E-07 | Yes |
| *NOTCH1* | -2.4 | 6.18E-06 | Yes |
| *NEO1* | -2.1 | 1.56E-05 | Yes |

To examine whether GLIS3 binding peaks were associated with transcriptionally active regulatory regions, we compared the GLIS3 genome browser tracks with ATAC-seq peaks retrieved from ChIP-atlas (ID-SRX11746108, human astrocytes). This comparison revealed a remarkable overlap between ATAC-seq peaks and GLIS3 binding loci in several astrocyte-associated genes, including *GFAP*, *FABP7*, *ID1*, and *HEY1* (Fig. 5) indicating that GLIS3 is bound to regions of accessible, transcriptionally active chromatin. These results are consistent with a role for GLIS3 in the direct transcriptional regulation of astrocyte gene expression.

**Figure 5.**
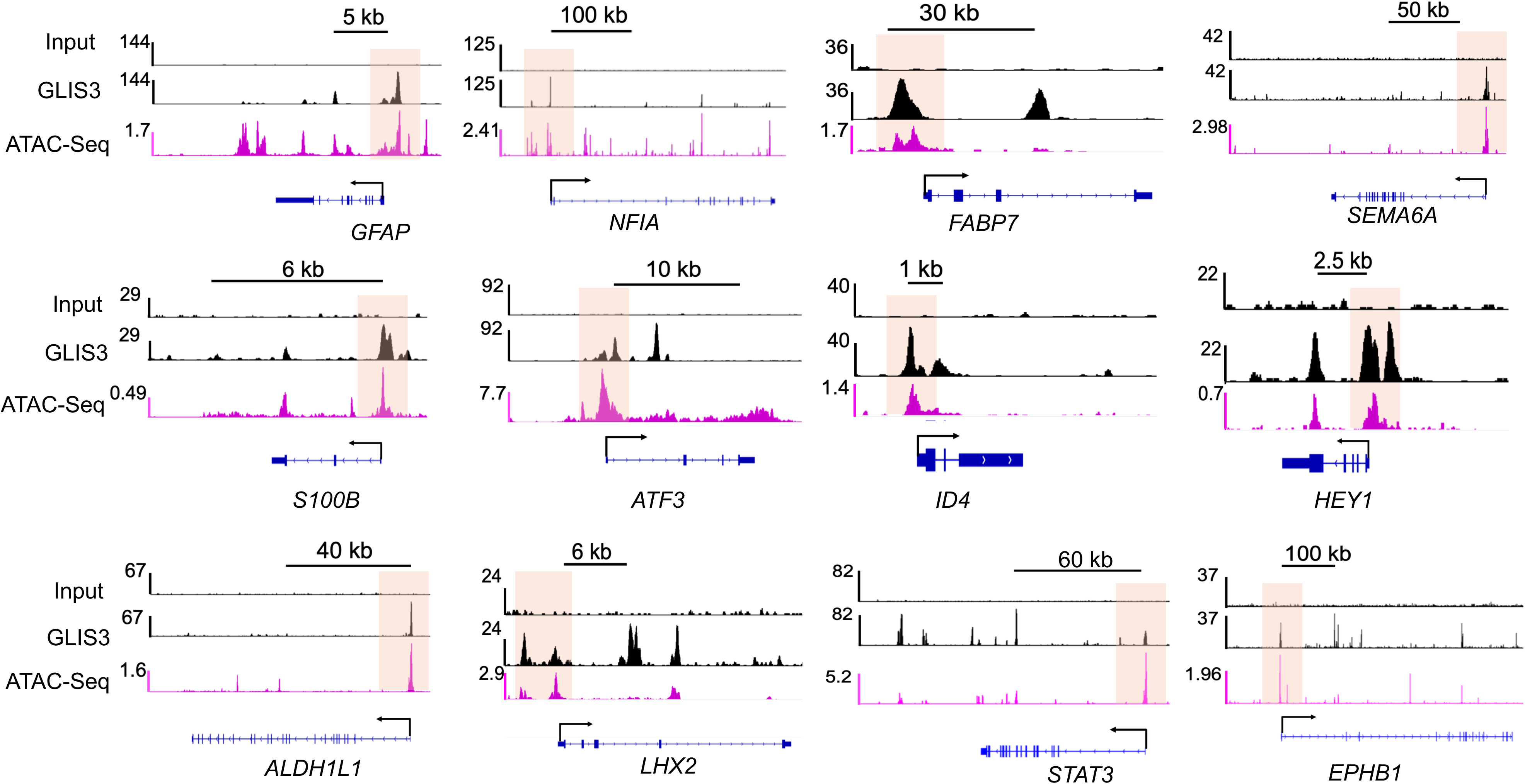
Genome browser tracks of several astrocyte GLIS3 target genes. Integrated genome viewer tracks of GLIS3 and ATAC-seq (human astrocytes SRX11746108 from ChIP-Atlas) showing location of GLIS3 binding and ATAC peaks. Input and GLIS3 represents the input control and GLIS3 ChIP data. ATAC peaks indicate regions of open chromatin.

### Exogenous GLIS3 expression promotes astrocyte differentiation in GLIS3^-/-^ NPCs

Since loss of GLIS3 function inhibited astrocyte differentiation, we questioned whether exogenous GLIS3 expression in *GLIS3*^-/-^ NPCs might restore astrocyte differentiation (Fig. 6A). To investigate this, we focused on the effect of exogenous GLIS3 on the expression *GFAP,* an established astrocyte marker and a GLIS3 target gene repressed in *GLIS3*^-/-^ cells (Fig. 3A; Table 1). As shown in Fig. 6B, exogenous GLIS3 expression induced *GFAP* mRNA to levels comparable to those in WT cells and this was accompanied by a dramatic increase in GFAP^+^ cells (Fig. 6C). The expression of several other astrocyte-related genes, such as *S100B, SLC1A1/3, ATF3,* and *NFIA,* were also significantly induced upon GLIS3 expression (Fig. 6D). These data together suggest that exogenous GLIS3 expression restores astrocyte differentiation in *GLIS3*^-/-^ NPCs consistent with the hypothesis that GLIS3 is a critical regulator of NPC differentiation into astrocytes. This was further supported by data showing that exogenous GLIS3 expression promoted astrocyte differentiation in WT NPCs (Fig. 6E). During the differentiation of WT NPCs into astrocytes *GFAP* was found to be minimally expressed at 7 days, moderately increased after 3 weeks, and highly expressed after 6 weeks of differentiation (Supplemental Fig. S7A). After 1-week, exogenous GLIS3 had little effect on *GFAP* expression (Supplemental Fig. S7B); however, after 4 weeks of astrocyte differentiation *GFAP* mRNA expression was about 8-fold higher in cells overexpressing GLIS3 compared to control cells (Fig. 6F). The higher expression of GFAP in overexpressing cells was supported by immune-blot and immunostaining for GFAP (Fig. 6G; Supplemental Fig. S7C). qPCR analysis further showed that the expression of the transcription factor *ATF3* and glutamate transporter *SLC1A2* was significantly higher (2-fold) in GLIS3 overexpressing cells compared to control cells. Although statistically not significant the expression of other astrocyte genes, such as *S100B* and *NFIA,* also shown an increasing trend (Fig. 6H). This suggests that expression of exogenous GLIS3 in NPCs in which GLIS3 is already highly expressed, has only a selective effect on enhancing the levels of astrocyte-associated genes.

**Figure 6.**
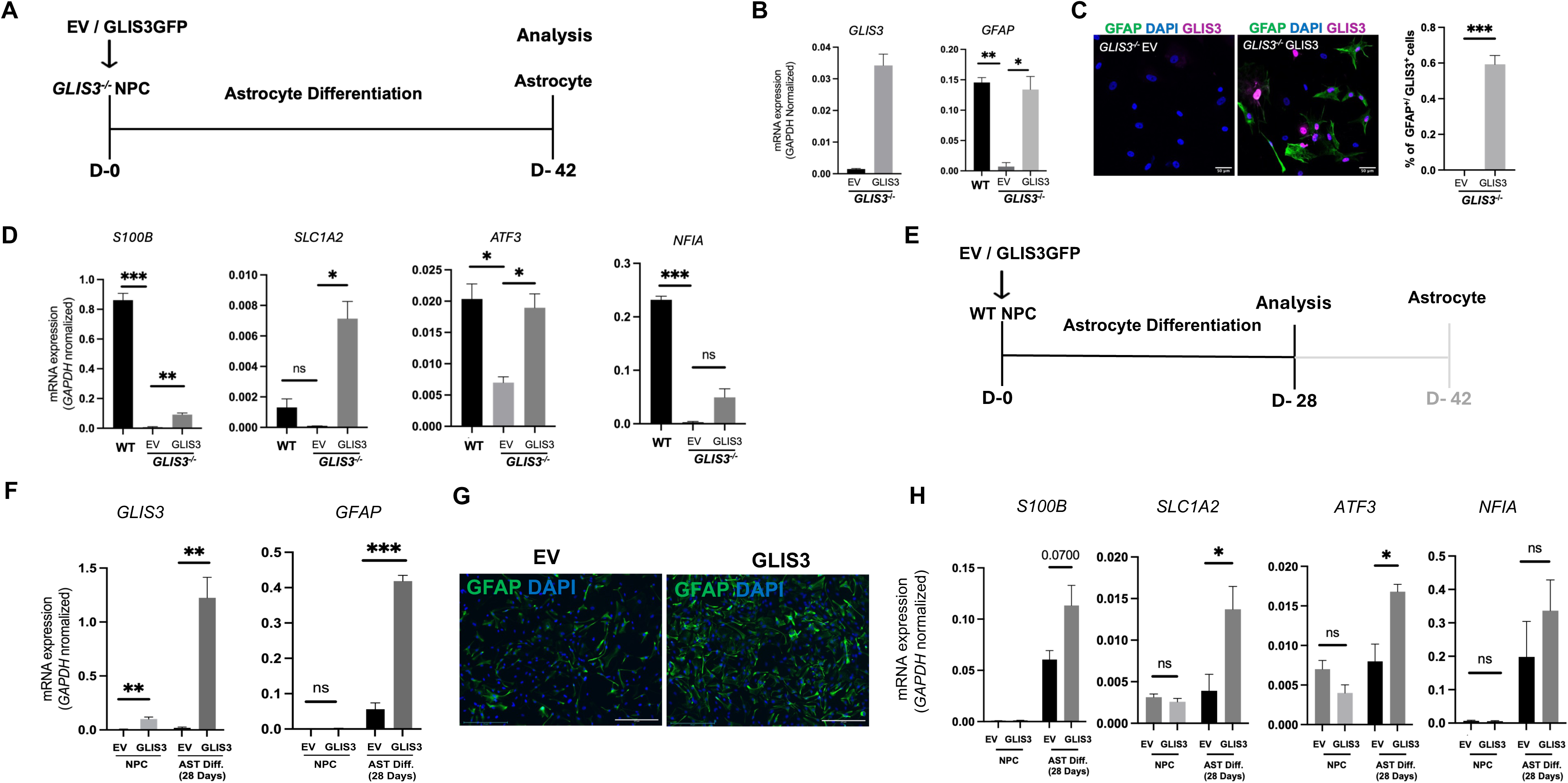
Exogenous GLIS3 expression promotes astrocyte differentiation in GLIS3^-/-^NPCs. (A) Schematic of the rescue protocol and timeline of *GLIS3 ^-/-^* NPC differentiation into astrocyte. (B) qPCR analysis of *GLIS3* and *GFAP* expression at end of astrocyte differentiation of WT and *GLIS3^-/-^* hNPCs infected with empty or GLIS3-GFP expressing lentivirus (n=3). (C) Co-immunostaining and quantification of GLIS3-GFP and GFAP in differentiated *GLIS3^-/-^* hNPCs infected with empty or GLIS3-GFP expressing lentivirus. (D) qPCR analysis of several genes astrocyte specific genes in *GLIS3^-/-^* hNPCs induced to differentiate into astrocytes with/without exogenous GLIS3 expression (n=3). (E) Schematic of protocol and timeline of GLIS3 overexpression on the differentiation of WT hNPCs into astrocytes. (F) qPCR analysis of *GLIS3* and *GFAP* levels in WT hNPCs infected with empty or GLIS3-GFP lentivirus at 28 days of astrocyte differentiation (n=3). (G) Immunofluorescence GFAP staining in WT hNPCs infected with empty or GLIS3-GFP lentivirus at 28 days of astrocyte differentiation. Scale bar 275 µm. (H) qPCR analysis of several astrocyte-related genes in WT hNPCs infected with empty or GLIS3-GFP lentivirus at 28 days of astrocyte differentiation (n=3). The data are presented as SEM values, with ∗ p<0.05; ∗∗ p<0.01; ∗∗∗ p<0.001; ns, not significant.

## Discussion

Differentiation of NPCs into astrocytes is a complex process that involves multiple signaling networks, such as the Notch and JAK-STAT pathway and interactions between various transcription factors, including SOX9, NFIA, and STAT3 (Miller and Gauthier 2007; Sloan and Barres 2014; Tyzack et al. 2016; Markey et al. 2023) . In this study, we describe a novel function of GLIS3 as a critical regulator of the differentiation of hNPCs into astrocytes and astrocyte gene expression. We show that GLIS3 is not only highly expressed in astrocytes at both the transcript and protein level but also that its expression progressively increases during differentiation of hESCs into NPCs and subsequently into astrocytes (Fig. 1). These observations suggested that GLIS3 might have a role in the regulation of this differentiation process and in the control of astrocyte function.

Study of the role of GLIS3 in the stepwise differentiation of hESCs into astrocytes showed that loss of GLIS3 function did not have any noticeable effect on the pluripotency of hESCs or their ability to differentiate into NPCs (Fig. 2). We previously reported that GLIS3 overexpression promotes the differentiation of hESCs into posterior NPCs in lieu of the default anterior pathway by inducing the expression of WNT3A, a strong posteriorizing factor (Jeon et al. 2019). As WT hESCs, *GLIS3*^-/-^hESCs differentiated mainly along the default anterior NPC lineage as indicated by the similar increase in the expression of the anterior marker genes, *OTX2* and *ZIC1*, while expression of the posterior marker genes, *GBX2* and *MNX1,* were not significantly induced in either WT or *GLIS3*^-/-^ hNPCs (Supplemental Fig. S3H), indicating that GLIS3 depletion had little impact on the NPC lineage. In contrast to the differentiation of hESCs into hNPCs, loss of GLIS3 function markedly impaired the differentiation of hNPCs into astrocytes. Expression of exogenous GLIS3 in *GLIS3*^-/-^ hNPCs was able to restore their ability to differentiate into astrocytes (Fig. 6) indicating that the repression of astrocyte differentiation was integral to the loss of GLIS3 function. Together, these findings demonstrate that loss of GLIS3 function impairs astrocyte differentiation consistent with a key regulatory role for GLIS3 in the differentiation of hNPC into astrocytes. Studies showing that *GLIS3* expression is repressed during *in vitro* differentiation of astrocytes into neurons (Rao et al. 2021) and suppressed during NPC differentiation into neurons (Tiwari et al. 2018), further support an astrocyte lineage-specific role for GLIS3. It has become increasingly apparent that astrocytes are heterogeneous and consist of various subpopulations that are in part determined by regional specificity (Gao et al. 2023; Kwon et al. 2025; Li et al. 2025; Williamson et al. 2025). Whether GLIS3 has different functions and regulates distinct genes in particular astrocyte subpopulations is an interesting question that warrants future investigation.

Since GLIS3 functions as a transcription factor, it is to be expected that it controls this differentiation and/or astrocyte function at least in part through its regulation of gene expression. Integrated transcriptome and cistrome analysis revealed that many genes involved in astrocyte differentiation and function were directly regulated by GLIS3 (Fig. 4). This included the transcription factor genes, *ATF3*, *SOX9*, *SOX2*, *LHX2*, and *NFIA*, and genes associated with astrocyte function, such as *GFAP*, *FABP7*, *S100B,* and *ALDH1L1.* Moreover, HOMER analysis identified, in addition to a consensus GLIS3 binding site, sequence motifs of several additional transcription factors, including members of the SOX, NFI, STAT, and AP-1 families, suggesting that they are located near GLIS3 binding peaks. Members of these families, including SOX9, SOX2, NFIA/B, STAT3, and ATF3, have been reported to have critical regulatory roles in astrocyte differentiation and/or astrocyte functions (Tiwari et al. 2018; Wang et al. 2022; Ma et al. 2024). Together, these findings suggest that GLIS3 regulate the transcription of a subset of astrocyte-related genes in coordination with these TFs. Although relatively little is known about the location of the binding peaks of these TFs in astrocytes, several studies have identified binding of several of these TFs to cis-regulatory regions in several astrocyte genes, some of which are located near GLIS3 binding peaks. For example, NFI-A/B, SOX9, and AP-1 bind *GFAP* within the same regulatory region as GLIS3 (Stolt et al. 2003; Gopalan et al. 2006a; Gopalan et al. 2006b). Similarly, study of Schwann and glioma cells identified binding of SOX and NFI members to cis-regulatory regions in *S100B* and *FABP7*, that are near the location of GLIS3 binding sites in astrocytes (Brun et al. 2009; Fujiwara et al. 2014). These data are consistent with our hypothesis that in astrocytes GLIS3 co-regulates certain genes in coordination with other TFs. Analysis of ATAC-Seq data from astrocytes revealed that GLIS3 binding sites are located within open chromatin, transcriptionally active regions (Fig. 5). Several studies have identified roles for *NFIA*, *SOX9, SOX8*, and *ATF3* in chromatin remodeling during astrogliogenesis (Kang et al. 2012; Tiwari et al. 2018; Lattke et al. 2021; Takouda et al. 2021). We previously reported that GLIS3 does not function as a pioneer factor in thyroid follicular cells and pancreatic beta cells (Scoville et al. 2020; Kang et al. 2023) but rather as a cooperating transcription factor (Balsalobre and Drouin 2022). Future studies need to determine whether this is also the case in astrocytes.

Our cistrome analyses identified several genes critical for astrocyte-associated functions as targets of GLIS3 transcriptional regulation. These include *GFAP*, the glutamate receptor *GRIA2*, the glutamate transporter *SLC2A1* (EEAT1), glutamate-ammonia ligase (*GLUL*), neuroligin-1 (*NLGL1*), neuroregulin1 (*NRG1*), potassium inwardly-rectifying channel, subfamily J, Member 10 (*KCNJ10*; Kir4.1), thrombospondin 3 (*THBS3*), semaphorin 6A (*SEMA6A*), and the GABA transporters *SLC6A1* (GAT-1) and *SLC6A11* (GAT-3), genes that are involved in the maintenance of glutamate and potassium homeostasis, and synaptic transmission(Mederos et al. 2018; Mahmoud et al. 2019; Wang et al. 2020; Liu et al. 2021). These observations suggest that GLIS3 may have a function beyond the regulation of astrocyte differentiation and has also a role in regulating astrocyte functions. Future studies are needed to determine whether loss of GLIS3 function GLIS3 affects astrocyte functions, synaptic transmission, maturation, and/or activation.

Using integrated transcriptomics and cistromics, our study provides the first comprehensive analysis of the role of GLIS3 in the differentiation of hNPCs into astrocytes and in the direct transcriptional regulation of genes with roles in astrocyte differentiation and function. Considering the association of *GLIS3* variants with several neurological disorders, such as Alzheimer’s and Parkinson’s disease, and the role astrocytes play in these pathologies, our study provides a valuable framework for exploring how GLIS3 might impact these diseases and may provide a new therapeutic target in the management of these disorders.

## Materials and Methods

### Cell culture

H9 human embryonic stem cells (hESCs)(WiCell, Madison, WI, USA) were grown in 6-well plates coated with 2.5% hESC-qualified Matrigel (Corning 354277) in mTeSR1 (Stemcell Technologies 85850) containing 10 μM ROCK inhibitor (Tocris Bioscience 1254). hESCs were passaged every 4–5 days, replated at a dilution of 1:6, and maintained in mTeSR1 in Matrigel-coated culture dishes.

### Mice

*Glis3*-EGFP mice (C57BL/6-Glis3<tm3(Glis3-EGFP)Amj>) expressing a GLIS3-EGFP fusion protein were described previously (Kang et al. 2017). Animal studies followed guidelines outlined by the NIH Guide for the Care and Use of Laboratory Animals and protocols were approved by the Institutional Animal Care and Use Committee at the NIEHS.

### GLIS3 knock-out hESCs

Three sgRNAs flanking the exon 4 region of *GLIS3* were designed (Supplemental Table S1) and cloned into CAS9 delivery plasmid PX459 V2.0 (Addgene; http://n2t.net/addgene:62988). Previous work has shown that deletion of any part of the DNA binding domain produces a non-functional protein (Beak et al. 2008). To generate GLIS3-knockout hESCs (*GLIS3*^-/-^ ESCs), sgRNA electroporation was performed using 4D-Nucleofector (Lonza Bioscience AAF-1003X) according to manufacturer’s recommendation. Briefly, H9 hESCs were harvested using accutase and 5 x 10^5^ cells electroporated with 10 µg plasmid. After electroporation, cells were plated in the presence of 10 µM Y27631 in mTeSR1 media. Individual knockout clones were isolated and sequenced to confirm the deletion. To address any observational variabilities in phenotypic characteristics of *GLIS3^-/-^* ESCs, two homozygous *GLIS3*^-/-^ESC clones, C20 (GLIS3^-/-^ -1) and C8 (GLIS3^-/-^ -2), containing different out of frame mutations, were chosen for further study (Supplemental Fig. S1).

### Astrocyte differentiation

All media/kits used for astrocyte differentiation were obtained from STEMCELL Technologies. hESC cells were differentiated into neural progenitor cells (NPCs) using a STEMdiff™ SMADi Neural Induction Kit (STEMCELL Technologies 08581), which were subsequently differentiated into astrocytes using an astrocyte differentiation and maturation kit (STEMCELL Technologies 100-0013, 100-0016), following the manufacturer’s protocol. Briefly, NPCs were seeded onto Matrigel pre-coated 6-well plates at a density of 2x10^5^ cells in NPC expansion medium. The next day, medium was replaced with astrocyte differentiation medium and renewed everyday with fresh medium for a week. Cells were subsequently dissociated with Accutase for 5 min, resuspended, and replated in Matrigel-coated wells and differentiation medium at a density of 2x10^5^ cells. Cells were maintained in differentiation medium for 3 weeks with intermittent passaging. After 3 weeks, cells were maintained in astrocyte maturation medium for three weeks and then analyzed for astrocyte differentiation markers.

### Lentivirus

Lentivirus expression plasmids: pLVX-mcherry-N1(control) and pLVX-mGlis3L-GFP (GLIS3) were used for transduction. Briefly, NPCs (3 x 10^5^ well) were transduced with lentivirus (∼9.2 X10^5^ titer/well) in a 6-well plate in astrocyte differentiation medium. The infection was performed twice in consecutive weeks beginning at the start of the differentiation.

### Primary murine astrocyte and microglia culture

Single cell suspensions were obtained from brains of PND3∼4 *Glis3*-EGFP mice using adult brain dissociation kit (Miltenyi Biotech 130-107-677). Primary astrocytes and microglia cells were enriched using ACSA2 and CD11B microbeads (Miltenyi Biotech 130-097-678,130-097-142) following the manufacturer’s protocol and grown in Poly-L-lysine (Sigma, Millipore P4707) coated dishes in glial cell culture media (Supplemental Table S2).

### RNA-sequencing and Analysis

Total RNA was extracted using PureLink^TM^ mini-RNA isolation kit (Thermo Fisher Scientific 12183018A). TruSeq Stranded mRNA kit and TruSeq RNA Library preparation kit (Illumina Inc., San Diego, CA, USA) were used to make libraries for RNA-Seq. RNA-seq data was generated as paired-end 76mers on a NextSeq 500 (Illumina, USA). Read pairs were then mapped to the hg38 reference assembly via STAR v2.5 (Dobin et al. 2013) with parameters “--outSAMattrIHstart 0 --outFilterType BySJout --alignSJoverhangMin 8 --limitBAMsortRAM 55000000000 --outSAMstrandField intronMotif --outFilterIntronMotifs RemoveNoncanonical”. Counts per gene were determined via featureCounts (Subread v1.5.0-p1) (Liao et al. 2014) with parameters “-s0 -Sfr -p”. Evaluated gene models are taken from the NCBI RefSeq Curated reference as downloaded from the UCSC Table Browser on July 10, 2020. Principal component and differential gene expression analyses were performed with DESeq2 v1.42.0 (Love et al. 2014); DEGs were defined based on an adjusted p-value threshold of 0.01. Gene expression heatmaps are generated by GraphPad using TMM-normalized CPM scores as reported by EdgeR v4.0.5 (Chen et al. 2025). Pathway analysis was performed using DAVID bioinformatics tool.

### Real-time qPCR

Total RNA was isolated from cells using a PureLink^TM^ mini-RNA isolation kit following the manufacturer’s protocol. On-column DNase treatment was performed using a DNase Kit (Invitrogen 12185010). The quantity of isolated RNA was measured using Nanodrop 2000 (Thermo Fisher Scientific ND-2000). 500 ng of total RNA was used for cDNA synthesis using a High-capacity cDNA conversion kit (Invitrogen 4374967) following the manufacturer’s protocol. 10 ng of cDNA was used as a template for Real-time qPCR analysis using SYBR green reagent (Invitrogen 4309155). *GAPDH* was used as an endogenous control for normalizing the target gene expression. The relative mRNA expression was calculated using the ddCt method. Sequences of all primers used for qPCR are provided in Supplemental Table S3.

### Immuno-cytochemistry

Briefly, cells were fixed in 4% paraformaldehyde (Electron Microscopy Sciences 15700) at room temperature (RT) for 10 min, washed twice with Dulbecco’s phosphate-buffered saline (DPBS), and then permeabilized with 0.2% Triton X-100 (Sigma Millipore T8787) for 15 min at RT. Cells were then washed twice with DPBS and blocked in DPBS containing 5% donkey serum for 1h. Cells were incubated with primary antibodies at 4°C in blocking buffer overnight and then washed three times with DPBS and subsequently incubated for 1h at RT with secondary antibodies and washed twice with DPBS. Antibodies are listed in Supplemental Table S4. Nuclei were identified by 4’, 6’-diamidino-2-phenylindole (DAPI)(Abcam ab228549). Fluorescence was observed with a Zeiss LSM-780 confocal microscope.

### ChIP-Seq sample preparation, library construction and analysis

GLIS3 expressing astrocytes (1 × 10^6^ cells) were cross-linked with 1% formaldehyde in PBS for 10 min at RT, and the reaction was quenched by the addition of 0.125 M of glycine at a final concentration for 10 min at RT as described previously (Jeon et al. 2019). Cells were washed twice with PBS, resuspended in 1ml of cold lysis buffer A (50 mM Hepes, pH7.5; 140 mM NaCl; 1 mM EDTA; 10% glycerol; 0.5% NP-40; 0.25%

Triton X-100; 1X Roche complete protease inhibitor cocktail (Sigma, Millipore 04693116001), and incubated for 10 min on ice. Cells were then pelleted and resuspended in 1 ml lysis buffer B (10 mM Tris-HCl, pH 8.0; 200 mM NaCl; 1 mM EDTA; 0.5 mM EGTA; 1X protease inhibitor cocktail). After 10 min on ice, cells were sonicated for 10 min in lysis buffer C (10 mM Tris-HCl, pH 8.0; 100 mM NaCl; 1 mM EDTA; 0.5 mM EGTA; 0.5% N-lauroylsarcosine; 1× protease inhibitor cocktail) using a Covaris sonicator to obtain 200–500 bp fragments. After removal of cell debris by centrifugation at 14,000 rpm, chromatin was incubated overnight at 4°C with HA-antibody and subsequently incubated for 4h with protein A/G-conjugated magnetic beads (Thermo Fischer Scientific 88802). The chromatin-bound beads were then washed, and reverse cross-linked as previously described (Heard et al. 2001). DNA sequencing library was generated with the ChIPed DNA using a NEXTflex Rapid DNA sequencing kit (Bio Scientific). Sequencing reads were obtained using a NextiSeq 500 Sequencing System (Illumina, USA). Raw reads were preprocessed with TrimGalore v0.6.7 [https://github.com/FelixKrueger/TrimGalore] (parameters: --quality 20 –illumina --stringency 5 --length 30) for quality and adapter trimming, followed by mapping to the hg38 reference assembly via Bowtie v2.1.0 (Langmead and Salzberg 2012) using the “--local” setting. Alignments were filtered for a minimum MAPQ score of 5, then duplicates were removed by MarkDuplicates.jar (Picard tool suite v1.110 (https://broadinstitute.github.io/picard/) with the REMOVE_DUPLICATES=TRUE option. Peak calling was carried out using the HOMER v4.11.1 (Heinz et al. 2010) ‘findPeaks’ function with parameters “-style factor -fdr 1e-5”. Peaks overlapping with hg38 blacklist regions were excluded from downstream analysis. Enriched motif analysis was performed by the HOMER v4.11.1 ‘findMotifsGenome.pl’ function using parameter “-size given”. For the purpose of the genomic context summary of ChIP-seq peaks, TSS proximal is defined as -1 kb relative to annotated transcriptional start site (TSS), Upstream is defined as -10 kb to -1 kb relative to TSS, gene body is TSS to transcription end site (TES), and Intergenic is all other genomic locations; these annotations are based on NCBI RefSeq Curated gene models. GLIS3 target genes are defined as those gene models for which an annotated TSS is within 5 kb of a ChIP-seq peak. For the visualization of genomic ChIP-seq signal (including heatmap, metaplot, and coverage tracks), mapped reads were extended to the estimated average fragment length (200bp) with the BEDTools v2.29.2 (Heinz et al. 2010) ‘slop’ function.

### Gene set enrichment and Gene Ontology analysis

Gene Set Enrichment Analysis (GSEA) software was used to analyze pre-ranked gene list from *GLIS3*^-/-^ vs WT RNA-Seq expression data. Pre-ranked gene lists were analyzed against GSEA’s Molecular Signature Database (MSigDB) gene sets (Mootha et al. 2003; Subramanian et al. 2005). Gene Ontology analysis was used identifying enrichment different bioprocess using DAVID bioinformatics tool (Huang da et al. 2009a; Huang da et al. 2009b).

### Statistical Analysis

All statistical analyses were performed using GraphPad Prism software version 11. Unpaired student t-test was performed for qPCR gene expression comparison among groups. One way ANOVA test was performed to analyze data between more than two groups. Biological replicates(n) represent independent experiments. *** indicates p <0.001, ** p <0.01, and * p <0.05, ns, not significant.

### Data availability statement

The ChIP-Seq and RNA-Seq data used in this study have been deposited in the Gene Expression Omnibus under accession numbers GSE319581 and GSE319582 respectively.

## Acknowledgments

We would like to acknowledge Laura Miller at the NIEHS for her assistance in producing the mice, the Epigenomics and DNA Sequencing Core at NIEHS for carrying out RNA-seq and ChIP-seq analysis, and members of the Cell Biology Group for their thoughtful comments throughout.

## Author contributions

TP: Conceptualization; experimental design; investigation; data analysis; writing; HSK: investigation; data analysis; KJ: Conceptualization; experimental design; investigation; data analysis; KYP: GLIS3^-/-^ hESC generation; SAG: Bioinformatic data analysis, figure preparation; AMJ: Conceptualization; writing; supervision.

